# Quantitative PCR for cannabis flower containing SARs-CoV-2

**DOI:** 10.1101/2020.06.06.112474

**Authors:** Kevin J. McKernan, Liam T. Kane, Yvonne Helbert

**Affiliations:** Medicinal Genomics Corporation

## Abstract

In January of 2020, COVID-19 became a worldwide pandemic. As many industries shutdown to comply with social distancing measures, the cannabis industry was deemed an essential business in most U.S. jurisdictions. Cannabis is manually farmed, trimmed and packaged. Employees and trimmers in cannabis grows have been reported to test qPCR positive for SARs-CoV-2 and as a result cannabis flower can be a potential inhaled SARs-CoV-2 fomite. Many of the comorbidities described in COVID-19 are also qualifying conditions for medical cannabis access. Bat guano has been identified as a rich source for novel coronavirus discovery and it is also a common fertilizer in the cannabis field. To better assess cannabis fomite risk we developed a SARs-CoV-2 quantitative PCR assay optimized to operate with a hemp flower background matrix. This assay was utilized to estimate the stability of gamma irradiated SARs-CoV-2 as a hemp flower fomite.

## Introduction

A HIV pandemic altered the landscape of medical cannabis legislation in California in 1996. Dennis Peron (a medical cannabis and HIV activist) authored Proposition 215 (The Compassionate Use Act) which legalized medical cannabis in California. Twenty years later, in 2016, Cannabis was still federally illegal in all states. Colorado, Washington and many other states have since legalized recreational use at the state level. In many ways, viruses like HIV, proliferated the medical cannabis initiatives and while these laws have been successful for many patients it is not without side effects.

The cannabis industry recently experienced a respiratory illness. E-cigarette vaping associated lung injury (EVALI) presented itself in 2019 as a “vape pen” related illness believed to be associated with Vitamin E Acetate, commonly used as a solvent for some clandestine vaporization pens (Bhat et al. 2020; Lal et al. 2020). The epidemic also involved a lower percentage of nicotine pens. 78% of patients were using untested black-market pens, some sourced from China (Control 2020). This disease resulted in 2,807 hospitalizations and 68 fatalities and began to tail off in November of 2019 just as the COVID-19 epidemic was beginning to spread. This disease, while not believed to be contagious, shares symptomatology with COVID-19. Respiratory stress, pneumonia and abnormal ground-glass opacities in chest CT scans are common in both diseases and it is unknown to what extent these diseases exacerbate each other (Henry et al. 2019).

In 2020, Cannabis dispensaries are finding themselves to be in an unexpected class of “essential” businesses during the COVID-19 pandemic. This is presumably due to the high number of patients served by cannabis dispensaries and the increased demand on scarce pandemic hospital capacity that could occur with a cannabis dispensary shutdown.

Annual cannabis use varies by country and is estimated to be over 15% in the United States as of 2017 (Breckenridge et al. 2019). Use rates have climbed since legalization most notably in the older population (Han and Palamar 2018). States that have medical cannabis access have shown decreased Medicare expenditure and decreased opiate overdoses (Bachhuber et al. 2014; Bradford and Bradford 2016a; Bradford and Bradford 2016b; Bradford and Bradford 2017; Bachhuber et al. 2018a; Bachhuber et al. 2018b; Wing et al. 2020).

86%-99% of COVID-19 deaths are comorbid with diabetes, hypertension, chronic obstructive pulmonary disorder (COPD), and cancer (Franki 2020; Guan et al. 2020). Many cannabis patients are treating these conditions directly or palliatively with cannabinoids (Pratt et al. 2019). While few states list all of these conditions as approved medical conditions for cannabinoids, many patients point to anecdotal cases of cannabinoid use for cancer and HIV symptom relief. A double-blind placebo controlled trial exists for cannabinoids and diabetes (Jadoon et al. 2016) and a randomized placebo controlled trial for HIV-associated sensory neuropathy (Abrams et al. 2007). Various other studies exist for cannabinoid use in many autoimmune and reactive oxygen species (ROS) related diseases (Russo 2018; Hill 2020). Cannabinoid use is also common in febrile epilepsy and has sparked concern with COVID-19 as it is unclear if cannabinoids will dull the immune response or limit cytokine storms.

Many of the complications of COVID-19 include cytokine storms, clots and strokes. There is preliminary evidence for the endocannabinoid system’s involvement in these conditions, but there are no double blind placebo controlled trials to guide any prescriptive use (Klein et al. 2000; Coetzee et al. 2007; Nagarkatti et al. 2009; Choi et al. 2019). The antiviral properties of cannabinoids have been studied in other RNA viruses like HCV; however, there is currently no data to support COVID-19 treatment with cannabinoids (Lowe et al. 2017). A few cannabis derived compounds (CannabisinA and Quercetin) have been computationally predicted to be potent SARs-CoV-2 main protease inhibitors, but these compounds are not currently tracked or quantitated in cannabis flower and no *in-vitro* data exists for these molecules applied to SARs-CoV-2 (Ngo 2020). Byrareddy *et. al.* suggest Cannabinoids could be used to reduced lung inflammation in SARs-CoV-2 patients (Byrareddy and Mohan 2020) and Wang et al report some cannabinoid extracts modulating ACE2 expression in COVID-19 gateway tissues (Wang 2020), but these are also preliminary findings and require further exploration to understand prescriptive use.

Comorbid COVID-19 patients are also prone to secondary infections from *Aspergillus fumigatus, Aspergillus flavus, Candida glabrata, Candida albicans, Klebsiella pneumoniae, Staphylococcus aureus* (Blaize et al. 2020; Chen et al. 2020; Duployez et al. 2020; Huttner et al. 2020; Koehler et al. 2020). Fifty percent of COVID patients die with secondary infections (Zhou et al. 2020). Many of these pathogens are cannabis endophytes or epiphytes and are the targets of existing cannabis testing regulation (Feeney and Punja 2015; McKernan et al. 2015; McKernan et al. 2016; Thompson et al. 2017; Punja et al. 2019; Vujanovic et al. 2020). The primary infection of SARs-CoV-2 is currently not being screened for in the cannabis supply chain.

In 2020, Cannabis growers also began to report employees testing positive for SARs-CoV-2 (Adams 2020). Concurrently, novel coronaviruses are being discovered in bat guano, a fertilizer commonly used in the cannabis industry (Valitutto et al. 2020). Given cannabis flower is an inhaled product that requires manual contact to grow and trim, we developed tools to detect SARs-CoV-2 RNA within the background cannabis matrix. We describe a modified CDC/EUA based quantitative PCR assay for detection of SARs-CoV-2 RNA that contains an additional cannabis internal control amplicon. This amplicon helps to address false negatives and false positives that might arise due to the complex cannabis flower matrix.

With prevalent cannabis use and surges in cannabis sales during lockdowns, it is safe to assume that Cannabinoid use and COVID-19 will intersect. There is no evidence to suggest cannabis is a replicative host for SARs-CoV-2, and the cannabis lighting and curing process may sterilize the plant from becoming a productive fomite; however, human contact, sneezing or even breathing after curing may still present elevated fomite risk for oral or vaporized routes of administration (Judson et al. 2020; Li et al. 2020; Rubens et al. 2020).

Studies investigating the viability of SARs-CoV-2 on various surfaces and indoor environments imply stability on certain surfaces for up to 7 days (van Doremalen et al. 2020). These studies utilized 1 million - 100,000 copies tissue-culture infectious dose [TCID_50_] per milliliter and witnessed a greater reduction in viability on copper surfaces than plastic surfaces. It is important to note that the viability dropped 3 orders of magnitude in 72 hours on plastic. It is unknown if dried cannabis (rich in antioxidants, 10-20% by weight THCA and CBDA) is more or less stable as a fomite, but the plastic cannabis is often stored in ranks as one of the more stable fomites.

To address fomite stability, gamma irradiated virus was inoculated into hemp flower and extracted and quantitated with qPCR across 7 days (Figure 12.)

Given this intersection, we believe it is imperative and preemptive to survey the cannabis supply chain for SARs-CoV-2 RNA to prevent other modes of disease transmission. Given the unexpected “essential” classification of cannabis dispensaries, we believe it is better to be proactive rather than reactive to potential risks that could change this status abruptly.

## Methods

We describe a single tube (Eppendorf) and 96 well plate format for processing samples. The 96 well method samples 5-fold less homogenate than the Eppendorf method due to volume restrictions on the 96 well plate (200 μL). This results in lower sensitivity for higher throughput methods but this smaller sampling volume can be compensated for by increasing the volume of the qPCR reaction or it can be addressed by using purifications of multiple 96 well preparations.

### DNA Purification

1-gram of USDA compliant (<0.3% (THC + 0.87 * THCA)) hemp flower (Spartan Hempworks, Massachusetts) is homogenized in 14.2mL of Tryptic Soy Broth (TSB) in a Nasco Whirlpak bag.

1mL of homogenate is removed and lysed with 50 μL MGC lysis buffer (Figure 1). This lysis buffer is known to effectively lyse open M13 Phage (Hawkins et al. 1997). Samples are vortexed and briefly spun with a table-top centrifuge. SARs-CoV-2 positive control DNA (IDT) was spiked in at various dilutions (200,000 copies, 20,000 copies, 2,000 copies, 200 copies).

**Figure 1.**
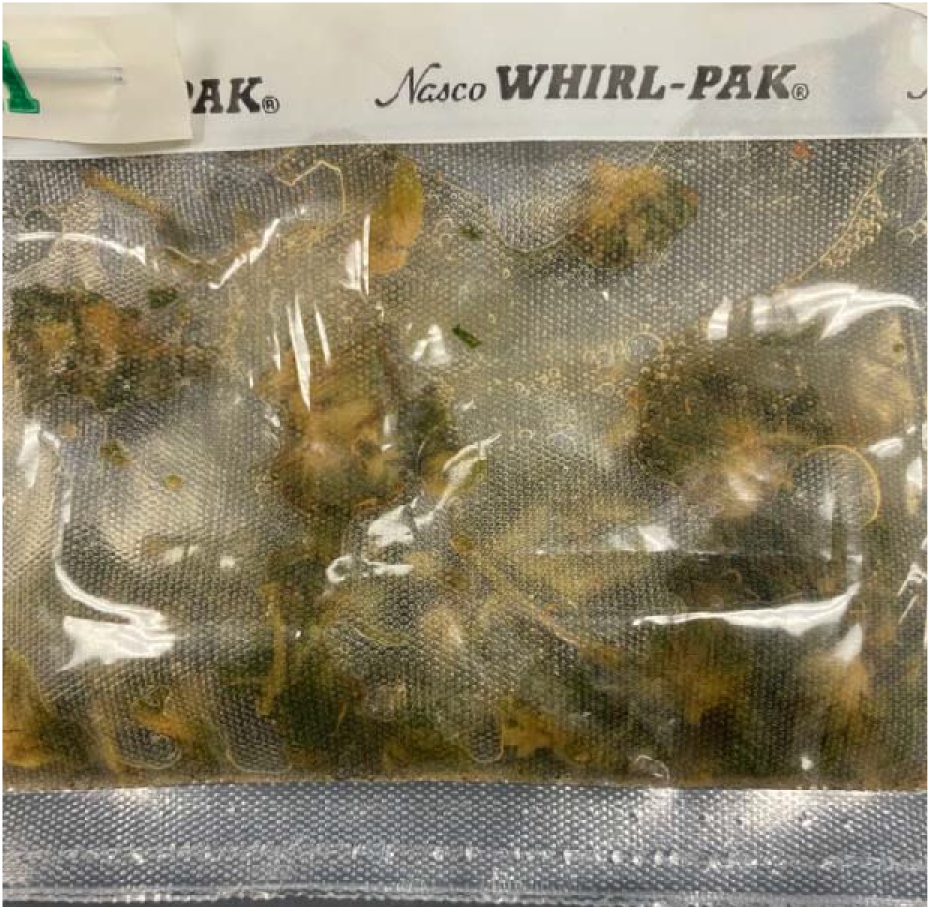
Hemp homogenized in 14.2mL TSB. 300 um mesh filters large plant debris from the sample within the WhirlPak bag. An additional spin step is required to pellet trichomes under 300 um in size.

### 96 well format

200 μL of lysate is purified using MGC nucleic acid SenSATIVAx purification system (SKU-420001). Samples are eluted in 25 μL of ddH20 and 8 μL of DNA is utilized in qPCR.

### Eppendorf format

1 mL of homogenate is removed from the WhirlPak bag from the back side of the 300 um mesh filter and placed into a 1.5 mL Eppendorf tube. 50 μL of MGC lysis buffer is added, vortexed briefly and spun down in a benchtop microcentrifuge (Figure 2). The supernatant is aspirated and placed into two separate Eppendorf tubes and 500 μL of MGC binding buffer is added, vortexed, incubated for 5 minutes and magnetically separated using a Magnetic Separation Rack (NEB #S1509S). Three 1 mL 70% ethanol washes are performed on the Eppendorf tubes. The resulting beads are dried for 20 minutes at RT and eluted in 25 μL ddH20 with a brief vortex. The 25 μL are combined into a single tube free of magnetic beads. For higher sensitivity, 21.5 μL of this eluent can be used in a 50 μL qPCR reaction.

**Figure 2.**
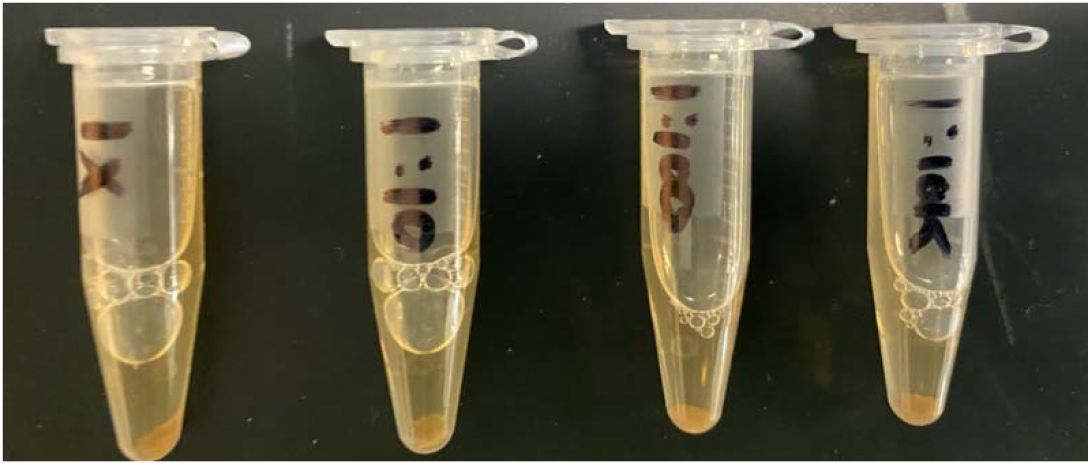
Eppendorf tubes after centrifugation shows visible signs of trichome pellets. The supernatant should contain lysed viral RNA.

Each format offers different throughput and different sampling. 20-100% of the 1 mL lysate is purified (96 well vs Eppendorf tubes). 96 well format is eluting in 25ul and utilizing 8.6ul in qPCR. This is results in 34% of the purified volume being placed into qPCR. 20% of the homogenate with 34% of the DNA sampled in qPCR results in 6.8% of the total 1 mL being assayed. This format offers lower sensitivity but higher throughput.

The Eppendorf format purifies the full 1 mL and utilizes up to 40% of the DNA with a 50 μL qPCR reaction. This approach is less automation friendly but offers over 6X higher sensitivity. Simply purifying more 96 wells per sample could increase the total homogenate volume assayed while still affording automated magnetic separation and washing. Variations of these two methods may find a more optimal compromise of cost, sensitivity and throughput. Failure to dry all of the ethanol out of the larger extractions can lead to failures that will be evident by the delay or lack of hemp internal control signal. Homogenized hemp samples usually produce 22-25 Cq.

### RNase viability proxy assay

Infectious SARs-CoV-2 must be handled in a bio-level 3 containment facility. SARs-CoV-2 DNA and RNA can be handled in bio-level 1 facilities and gamma irradiated virus requires bio-level 2 facilities. This places constraints on measuring viral “viability” that is traditionally assayed by counting plaques on Vero cell lawns. To estimate this without utilizing live cultures that replicate the virus, we emulated a live-dead assay from the microbial testing field that utilizes DNase to digest free circulating DNA while leaving genomic DNA inside a cell membrane intact (Medicinal Genomics part #420145). We hypothesized that RNases could achieve the same result with SARs-CoV-2 as RNase A shouldn’t traverse the nuclease protective envelope protein and only digest uncoated viral RNA.

To assay this, 100 μL of TSB cannabis homogenate is aspirated and split into 50 μL RNase negative and 50 μL RNase positive fractions. 4 μL of 20 mg/mL RNase A (New England Biolabs #T3018L) is added to the RNase positive aliquot of non-lysed TSB from 1-gram cannabis flower homogenate. The digestion is held at 37 °C for 30 minutes. Samples are purified using 50 μL of SenSATIVAx, magnetic separation, three 70% ethanol washes, and eluted in 25 μL ul of ddH20 and 2 μL of RNase blocker. Both RNase + and RNase-samples are compared in qPCR.

### Quantitative PCR

#### 20ul Reaction

10 μL MGC RT-qPCR master mix (Medicinal Genomics PathoSEEK^®^ COVID-19 Cannabis RT-qPCR Assay Kit #420209) 0.2 μL MGC primer

0.2 μL DTT

<u>1 μL RNase blocker</u>

11.4 μL Total

+

<u>8.6 μL DNA</u>

20 μL Total Reaction volume

#### 50ul Reaction

25 μL MGC RT-qPCR master mix

0.5 μL MGC Primer probe.

0.5 μL DTT

<u>2.5 μL RNase blocker</u>

28.5 μL Total

+

<u>21.5 μL DNA</u>

50 μL Total Reaction volume

PCR conditions.

Step 1) 50 °C for 15 minutes (RT step)

Step 2) 95 °C for 2 minutes (Denaturization)

Step 3) 95 °C for 5 seconds

Step 4) 55 °C for 45 seconds

Step 5) Goto Step 3, 49 more times.

## Results

DNA purification of cannabis homogenate (Figure 1) was explored in 96 well and single tube (1.5 mL Eppendorf) format (Figure 2 and 3). The N2 amplicons multiplexed with Cannabis internal control amplicons showed linearity from 500,000 copies down to 5 copies with SARs-CoV-2 DNA positive controls (Figure 4). The addition of the cannabis internal control assists in identifying DNA purification or sampling related false negatives (Figure 5,6). When assayed in the presence of subsampled hemp background this sensitivity decays from 5 copies to 50 copies (Figure 7,8) presumably due to competition with the multiplexed internal control. The N2 amplicon performs better than N1 amplicon (data not shown) with the added multiplexed internal control. Gradient qPCR and altered primer concentrations were explored to rescue the multiplexed N1-internal hemp control amplicon without success. Assay robustness was explored with varying lysis buffer concentrations (Figure 9) to ensure full lysis of the virus.

**Figure 3.**
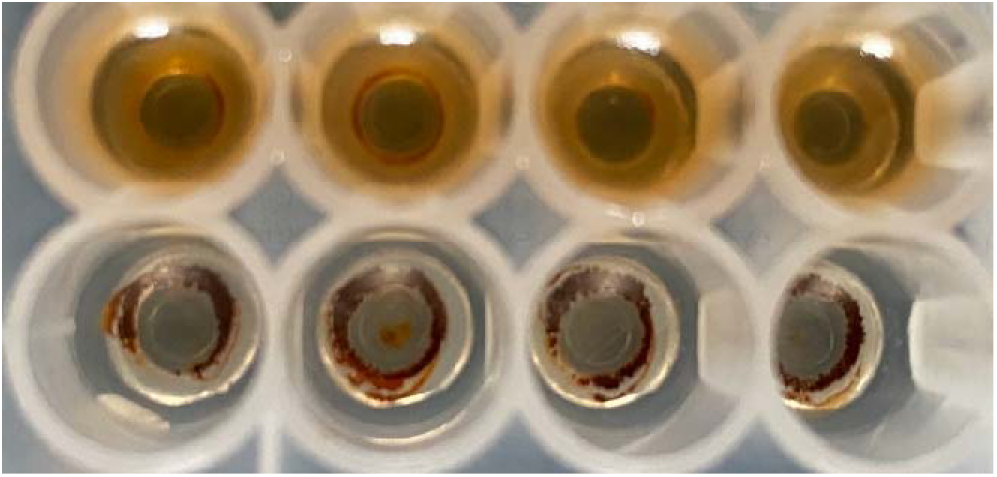
96 well format using magnetic isolation of RNA enables sample concentration prior to qPCR. 200 μL of homogenate is concentrated into 25 μL of eluant.

**Figure 4:**
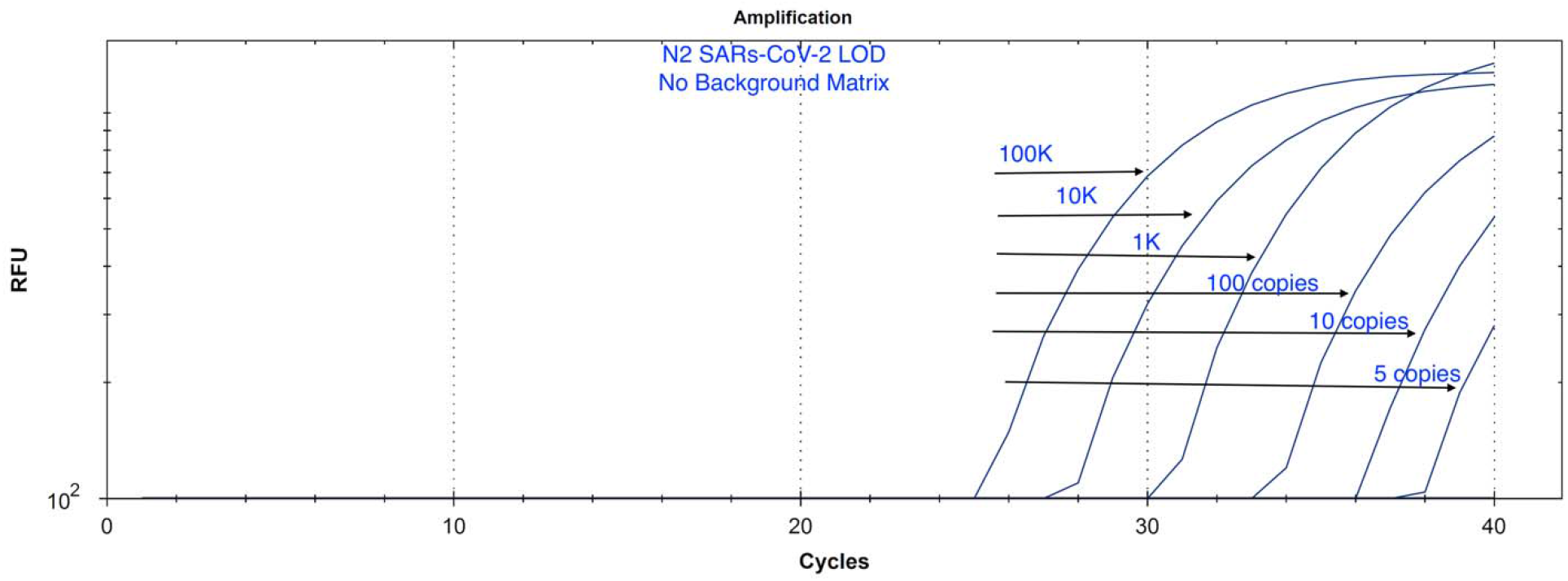
Assay LOD without Hemp Matrix. 10-fold serial dilutions of SARs-CoV-2 plasmid DNA (100,000, 10,000, 1,000, 100, 10, 5 copies of positive control). Lowest dilution equates to 80 copies in the original 1mL sample. The LOD of this assay is limited by sampling bias and would benefit from an internal PCR control to ensure the DNA purification and sampling was done correctly. As a result, we targeted a Cannabis DNA sequence that should amplify at higher copy number in every sample.

**Figure 5.**
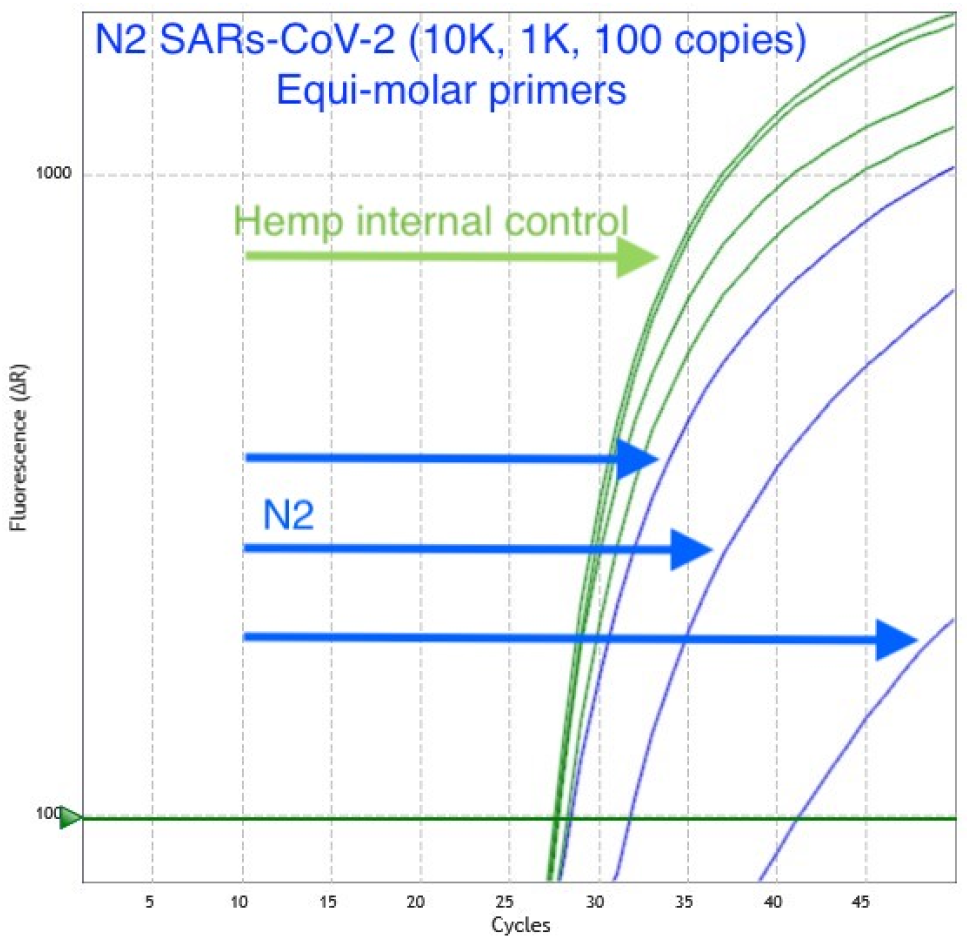
Assay LOD with Hemp Matrix. Serial dilution of SARs-CoV-2 positive control into hemp matrix. qPCR is detected on an Agilent Aria in FAM and HEX. BioRad CFX96 thermocycler was also tested. Hemp signal (Green/HEX) is relatively constant while SARs-CoV-2 amplicon N2 Cq (Blue/FAM) decays with 10-fold dilutions.

**Figure 6:**
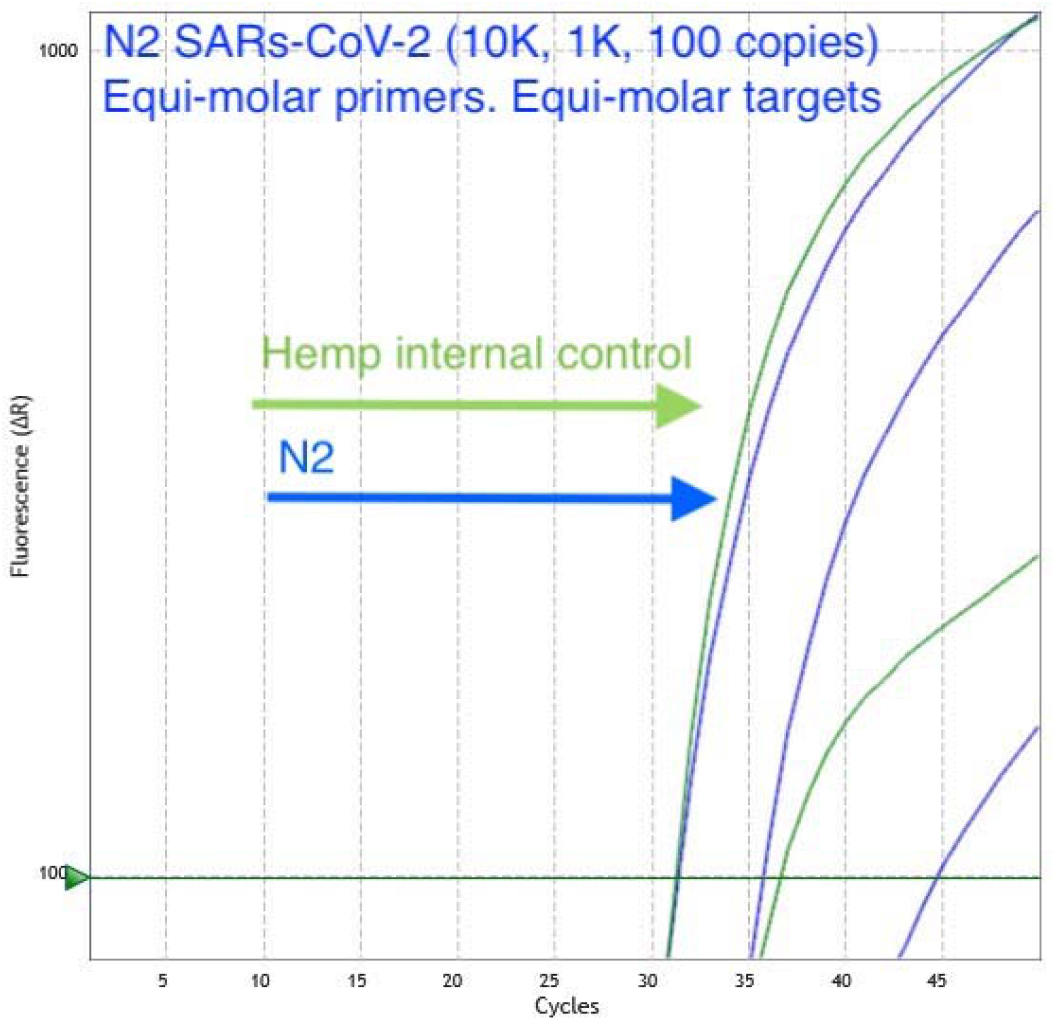
Assay LOD with Hemp Matrix. Serial dilution of both hemp and SARs-CoV-2 positive control demonstrates the N2 assay remains more sensitive than the hemp internal control as we approach the LOD. qPCR is detected on an Agilent Aria in FAM and HEX fluorescent channels. Hemp signal (Green/HEX) is relatively constant while SARs-CoV-2 N2 amplicon Cq (Blue/FAM) decays with 10-fold dilutions. This encouraged the reduction of the cannabis control amplicon primer concentrations to achieve even amplification efficiencies at low target concentrations with high cannabis background DNA.

**Figure 7.**
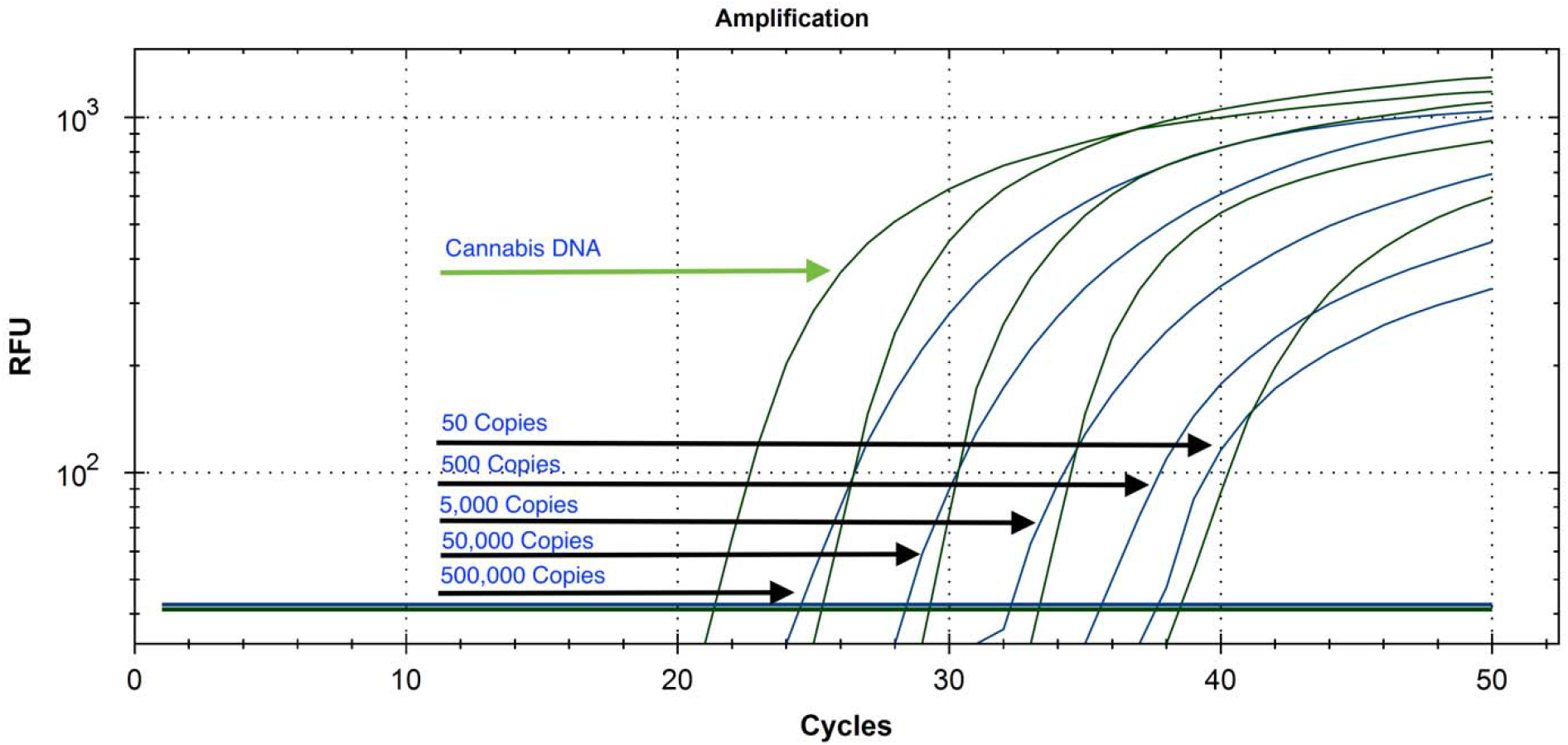
Serial dilution of plasmid SARs-CoV-2 DNA from 500,000 copies to 50 copies with cannabis internal control multiplexed on a Bio-Rad CFX-96.

**Figure 8.**
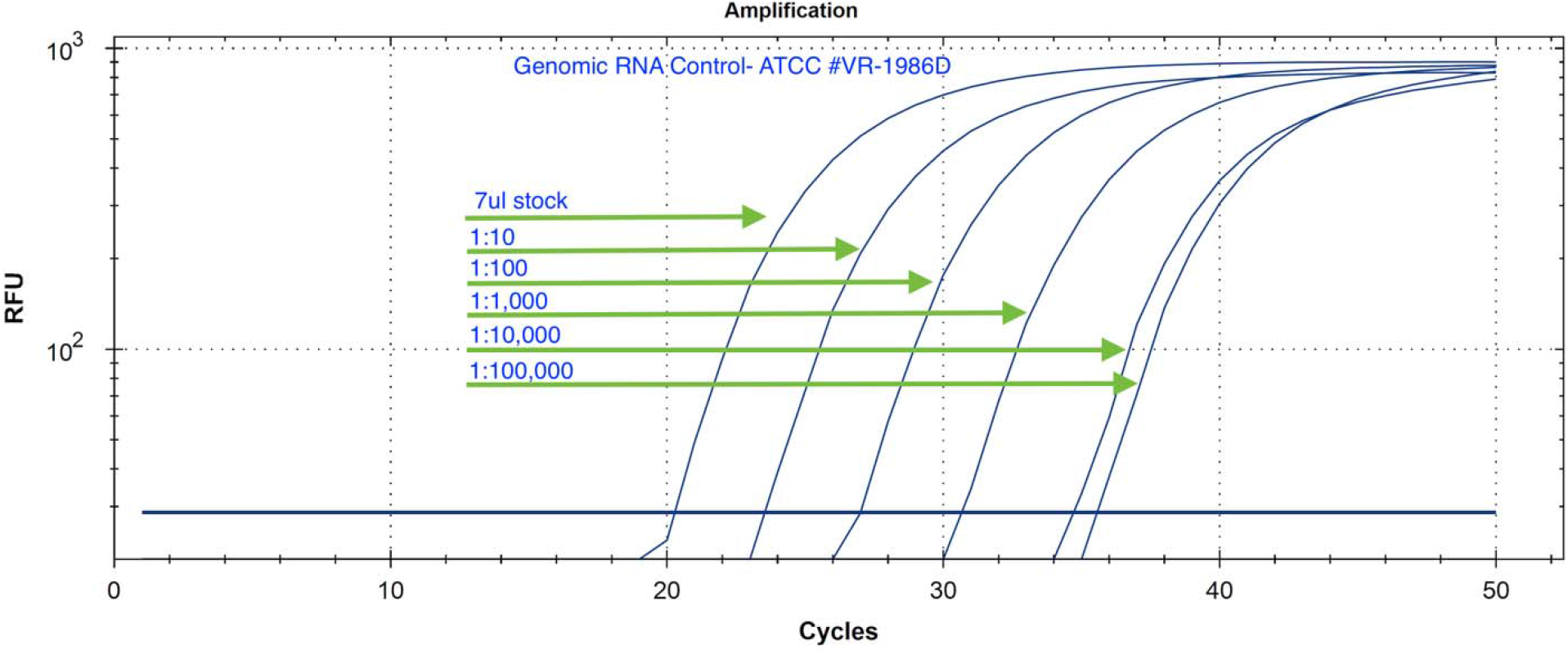
Serial Dilution of Genomic RNA isolated from a patient in Washington state in ddH20 (ATCC # VR-1986D). The initial concentration is from the manufacturer is not known but demonstrates linear dilution over 5 logs. No amplification is seen with the negative controls in 50 cycles.

**Figure 9.**
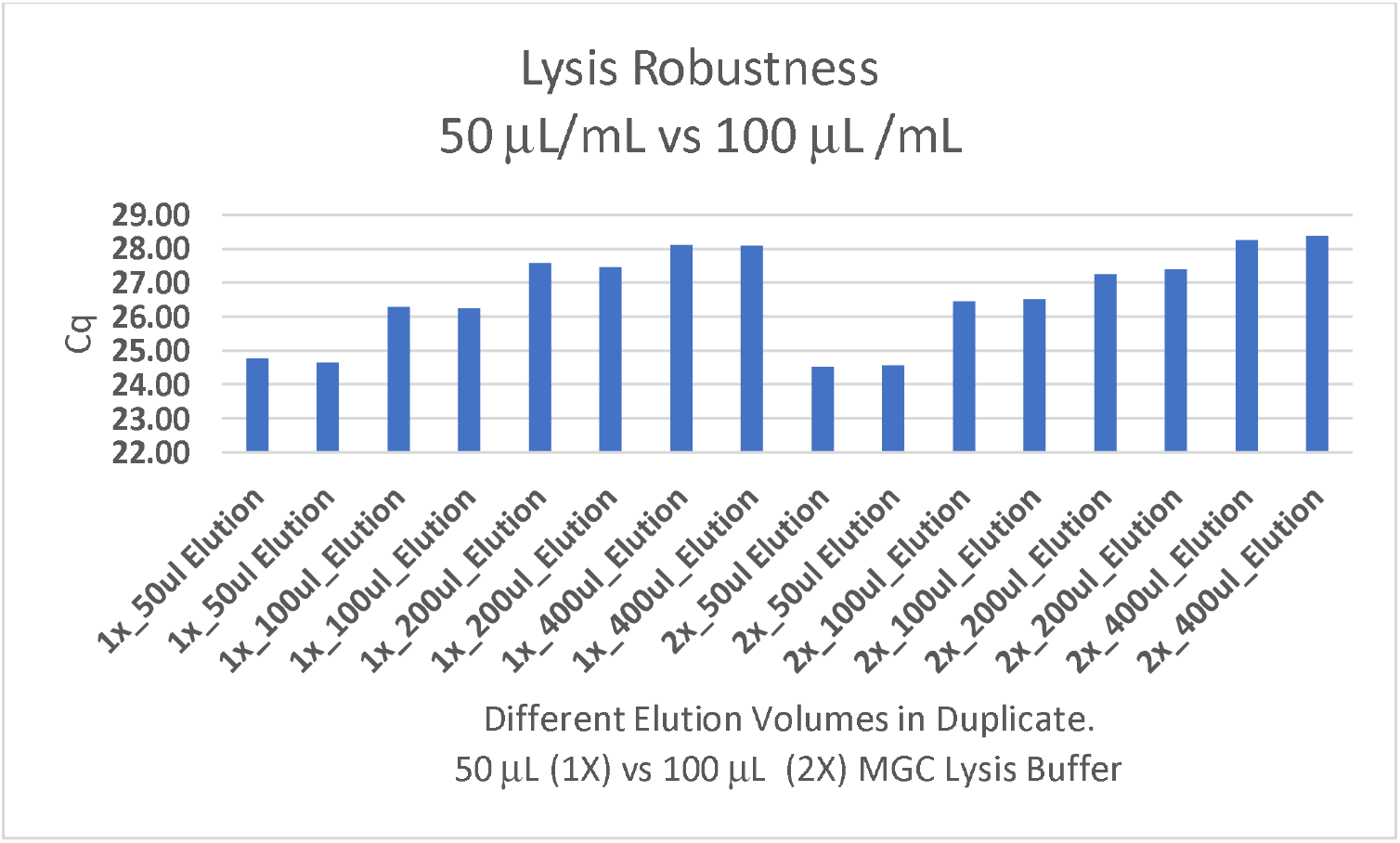
Lysis buffer robustness assessment. 50 μL/mL and 100 μL/mL of MGC lysis buffer applied to 1 mL of hemp homogenate was explored across 4 different elution volumes to optimize sample concentration and its impact on sensitivity. Cq’s reflect cannabis internal control target (HEX). Further reduction in elution volume may increase sensitivity.

Two additional SARs-CoV-2 positive controls were tested. A RNA whole genome control and a gamma irradiated attenuated virus were tested (ATCC-1986D & BEI Resources-52267) (Figure 10). The gamma irradiated reagent was deposited by the Centers for Disease Control and Prevention and was obtained through BEI Resources, NIAID, NIH: SARS-Related Coronavirus 2, Isolate USA-WA1/2020, Gamma-Irradiated, NR-52287. Both controls showed equivalent amplification to plasmid DNA based controls and the lysis buffer was confirmed to be effective at lysing SARs-CoV-2 virus particles.

**Figure 10.**
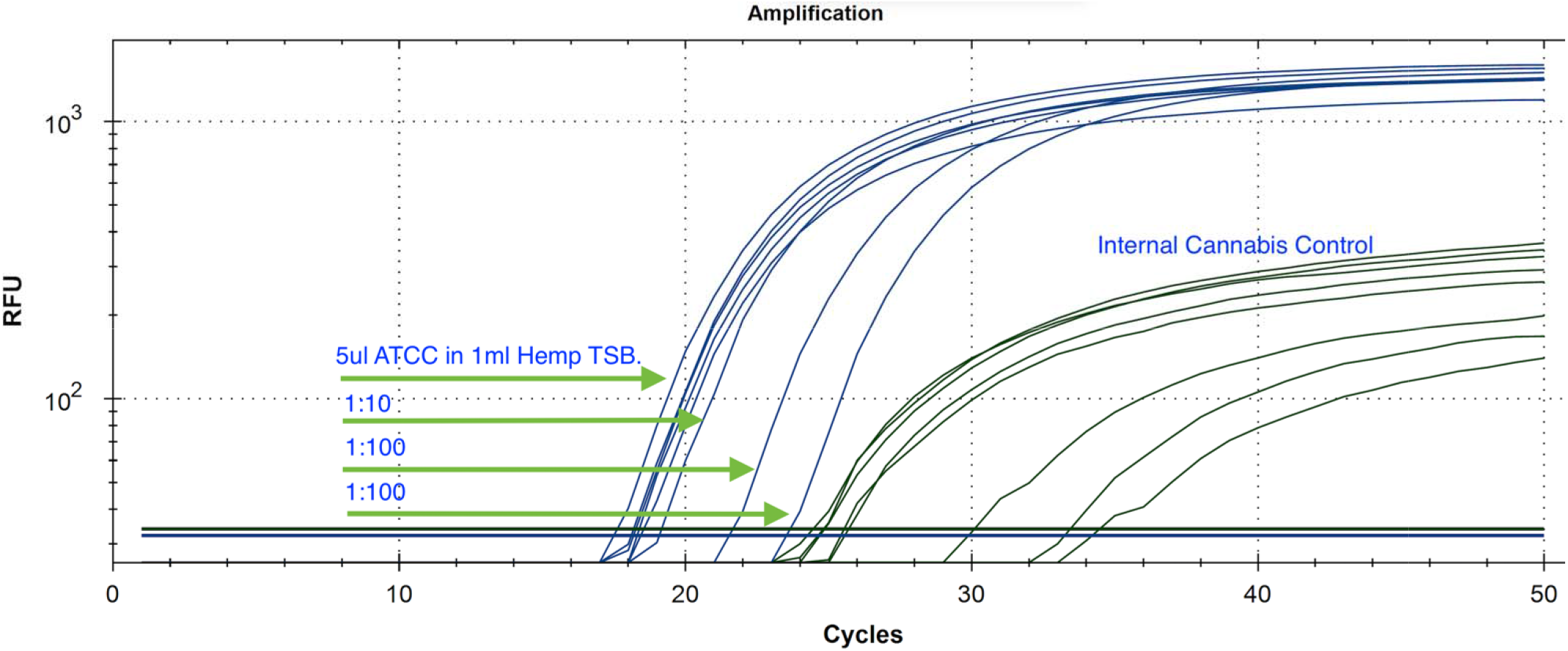
Gamma irradiated SARs-CoV-2 virus particle was diluted into 1mL cannabis homogenate and assayed with N2 amplicon and Cannabis Internal control targets (BEI Resources NR-52287). The gamma irradiated reagent was deposited by the Centers for Disease Control and Prevention and was obtained through BEI Resources, NIAID, NIH: SARS-Related Coronavirus 2, Isolate USA-WA1/2020, Gamma-Irradiated, NR-52287.

The assay sensitivity in cannabis matrix is affected by subsampling. One seventh to one tenth of the homogenate is purified (10-14 mL homogenate) and 30%-40% of the purified DNA is assayed in qPCR without the use of an enrichment step. This is equivalent to FDA approved assay from Roche on nasopharyngeal swabs where 1/5^th^ of the homogenate (0.2 mL/1 mL)is purified and 25% of the DNA assayed in qPCR (Diagnostic 2020).

In most jurisdictions a 1-gram sample of dried cannabis flower is required for pathogen testing. It is difficult to homogenize the entire 1-gram volume of cannabis (Figure 11) into a PCR assay. The 10 mL to 14.2 mL volumes assayed for cannabis are much larger than the 1 mL nasopharyngeal swabs used in human SARs-CoV-2 testing. DNA purification that elutes in a smaller volume than the homogenate concentrates the sample and can compensate for some of these expanded volumes required for cannabis sampling. Likewise, viral enrichment may overcome the sampling bias, but viral enrichment requires culturing the virus in the presence of Vero cells (BL3) and may not be advised given the risk to the testing lab employees (Colson et al. 2020; La Scola et al. 2020). Differential centrifugation techniques or antibody enrichment may improve virion sampling, but these may require itemization of packaged versus inactive RNA with RNase A as described.

**Figure 11.**
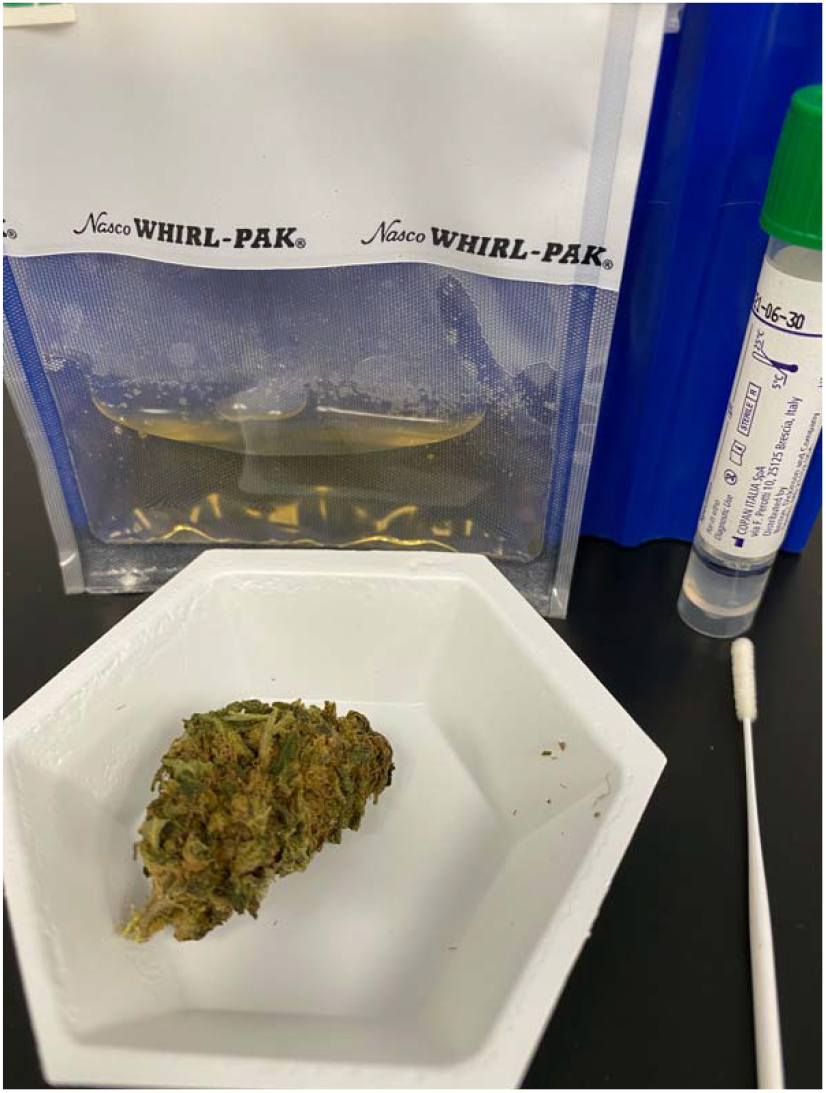
An example of sampling volumes utilized with Nasopharyngeal swabs (right) compared to Cannabis sampling techniques (left). Swabs are submerged in 1mL of lysis buffer (green cap tube), and 1/5^th^ of that sample (200 μL) is purified and eluted in 50μL. 10 μL of this RNA goes into qPCR. As a result, there is a 25-fold subsampling with Swab tests according to FDA/EUA approved Roche protocol (https://www.fda.gov/media/136231/download). 1-gram of cannabis doesn’t homogenize into 1mL of lysis buffer. As a result, Cannabis flower is homogenized in 10-14 mL of TSB and 1 mL is purified (7-10%) and eluted in 40 μL. If 40% of this DNA (20 μL of 50 μL) is placed into a 50 μL qPCR reaction a 25-fold subsampling can be achieved and presents the same sampling bias as the approved clinical tests being used to diagnose patients.

Gamma-irradiated SARs-CoV-2 is stable on non-sterile cannabis flowers for at least 7 days with the inoculations used in this study (Figure 12). The Cq shift from Day 0 to Day 7 represents a 120 fold decrease in RNA. This study only evaluated irradiated virus. Further work (BL3) is required to assess infectious versus non-infectious virus on cannabis. Hemp microbial regulations in the US are less stringent than medical cannabis microbial regulations. The high background total aerobic count on these samples may alter viability of the virus and should be repeated with samples sterilized more aggressively for medical cannabis use. Gamma-irradiated virus may have a shorter shelf life than infective virus and the infectability of gamma-irradiated virus is intentionally attenuated.

**Figure 12.**
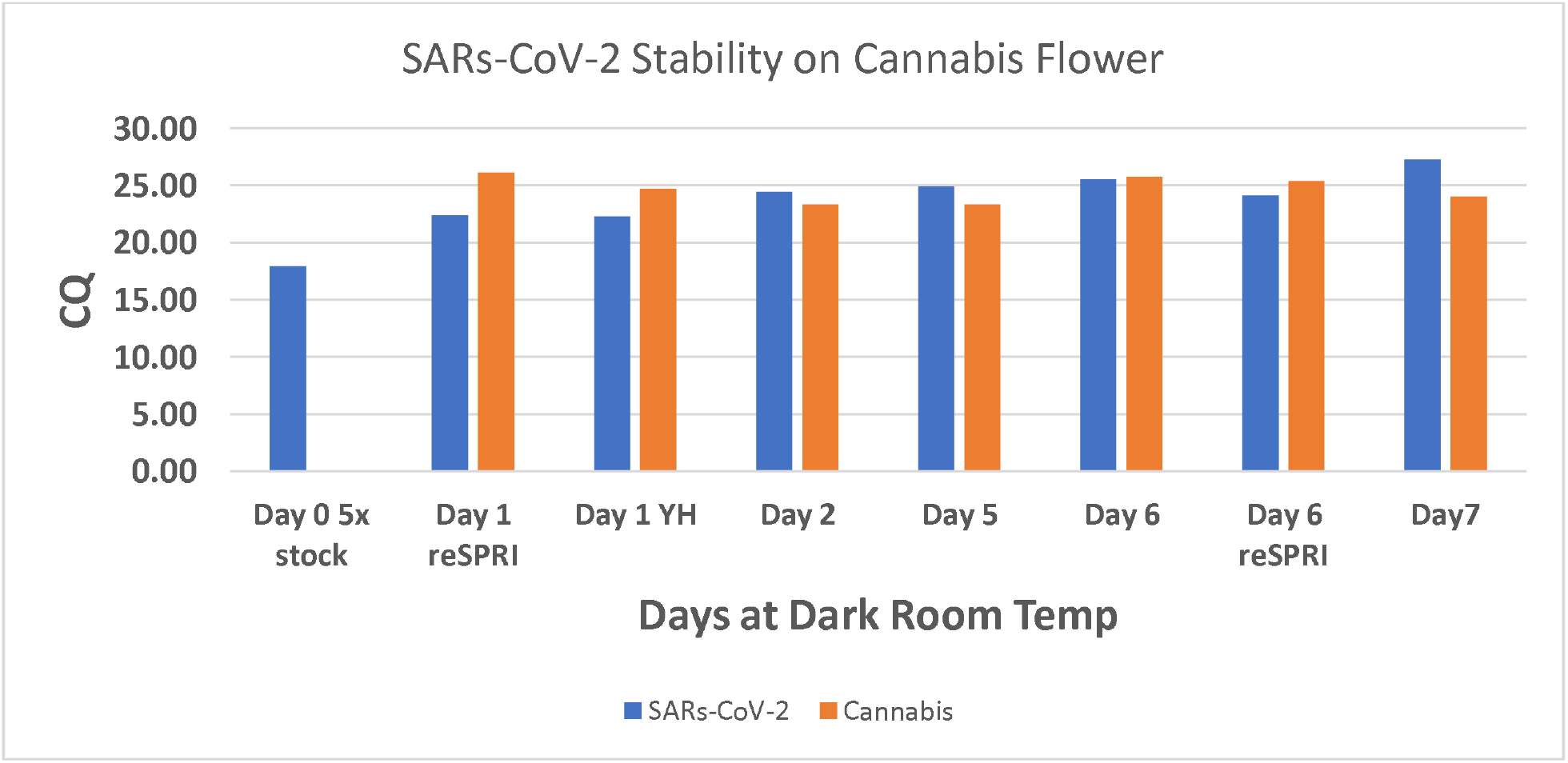
10 μL of a Gamma Irradiated SARs-CoV-2 (BEI Resources) was inoculated onto seven 1-gram non-sterile hemp flowers. Each sample was purified and assessed by qPCR run daily to quantitate RNA stability. Hemp flower was also assessed for Total Aerobic Count via qPCR and tested at over 1 million CFU/gram (data not shown).

To assess the “live-dead” status of the virus we RNase treated the viral RNA genome, viral DNA genome, and the gamma-irradiated genome to ascertain if the gamma-irradiated viral RNA still had protection from exogenous RNases (Figure 13). As expected, the RNase had no impact on the gDNA but showed considerable decay in qPCR Cq of the gamma-irradiated virus suggesting the viral envelop is not fully intact. The RNA genome was fully degraded suggesting the gamma-irradiated virus has a minimal amount of RNase protected genome. A Cq to PFU calibration with SARs-CoV-2 inoculated Vero cells (BL3 laboratory) is required to draw conclusive estimates of viral infectability on cannabis.

**Figure 13.**
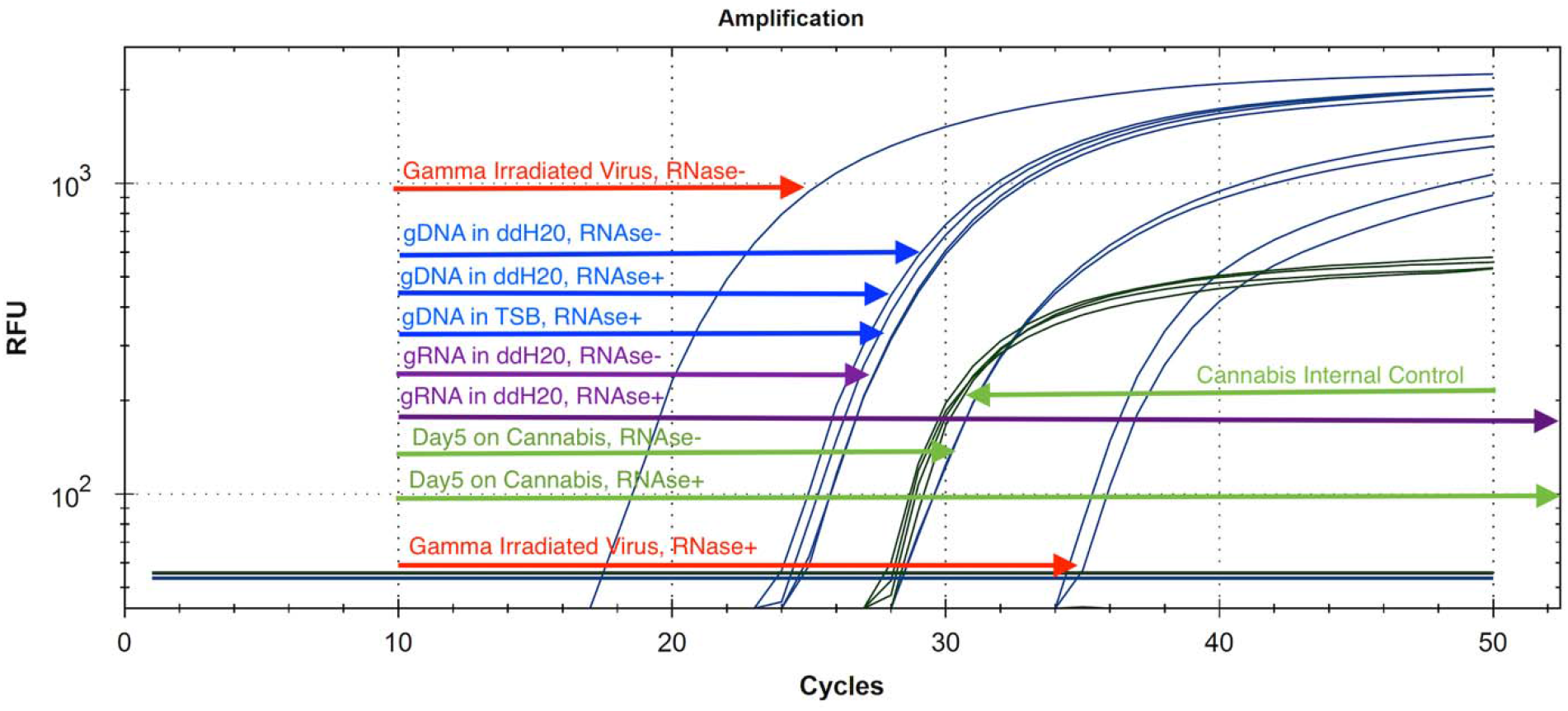
The use of RNase A as a proxy for viable virus versus non-viable virus. RNase should not penetrate the nuclease resistant viral envelope and only digest free circulating RNA. After purification of the RNase digested samples (37°C, 30 minutes), samples are eluted in the qPCR RNase Blocker. Gamma irradiated virus (Red) is less digestable than pure genomic RNA (Purple). Viral gDNA remains resistant to RNase digestion (Blue). Gamma irradiated virus spiked onto cannabis flower is fully digestable (Green) by RNase suggesting non-infectious virus by Day 5 at Room Temperature storage. Non-irradiated virus is expected to last much longer on cannabis.

## Discussion

The infective dose of SARs-CoV-2 virus particles is not currently known; however, superspreader events are believed to present higher viral load and result in more patients that progress to symptomatic disease (Wong et al. 2015). Aerosol and surface stability of SARs-CoV-2 has been measured on various surfaces (Doremalen). These studies evaluated thousands to millions of copies of the virus. Many described human SARs-CoV-2 LAMP and qPCR tests report similar sensitivity in the 10-100 copy range (Lau et al. 2003; Bruce 2020; Park 2020; Schmid-Burgk 2020). Digital PCR technologies have been proposed to enhance the sensitivity of qPCR as the sampling approaches single molecule levels, but these methods add significant costs and may not address the fundamental problems of low volume PCR reactions surveying and subsampling larger volumes of biological fluids (Dong 2020).

One limitation to this study is the use of cured cannabis samples. The polymerase utilized in this study has reverse transcriptase (RT) activity and will amplify both DNA and RNA. Non cured samples are likely to have higher RNA levels which may favor amplification of the cannabis internal control over the viral target. Experiments with DNAse treatment of these hemp samples demonstrated major shifts (5-8 Cq shift) in Cq suggesting very little contribution from Cannabis RNA but this may need further investigation for testing un-cured cannabis or different cannabis tissues.

To date, SARs-CoV-2 has never been detected on cannabis and cannabis is not believed to be a corona virus replicative host. It is possible for manually curated cannabis flower to become a fomite. It is doubtful virions will survive extraction or pyrolysis of cannabis flower but vaporization of cannabis flower and cannabis pre-rolls are documented microbial vectors (Remington et al. 2015; Shapiro Bb Md et al. 2018). Sterilization of SARs-CoV-2 requires 65 °C treatment for 30 minutes (ATCC part # 1986HK). Cannabis vaporization is usually performed over 130 °C but with a few seconds of duration. It is not yet known if this process will properly sterilize cannabis of SARs-CoV-2, but it has failed to sterilize cannabis of *Aspergillus* spores and resulted in clinical cases of Cannabis derived aspergillosis.

While chances of cannabis supply chain contamination with SARs-CoV-2 are remote, the RNA of the virus is stable on unsterilized cannabis flower for over 7 days and may inform personal protective gear protocols or sterilization techniques to ensure a virus free production process. A published SARs-CoV-2 surveillance method can greatly aid community preparedness and supply chain management. The sensitivity of this test is in line with the sensitivities being reported in nasopharyngeal swabs and other early SARs-CoV-2 amplification assays. More work is required to validate this on larger sample numbers and more diverse cannabis cultivars. This study evaluated DNA, RNA and gamma irradiated positive controls and primer sequences published in most EUA approved kits. Since there is no current evidence of SARs-CoV-2 detected on Cannabis, this study should not be interpreted as a justification to limit patient access to cannabis. The current COVID epidemic is a reminder of the importance of pre-emptive testing readiness and we believe these methods can contribute towards such goals.

## Author Contributions

KJM-Authored the manuscript, performed lab and analysis work

LTK-performed qPCR and DNA isolation work

YH-Performed qPCR and reagent procurement

## Acknowledgements

We want to thank Sherman Hom for thoughtful commentary on the manuscript.

